# SARS-CoV-2 Spike protein suppresses CTL-mediated killing by inhibiting immune synapse assembly

**DOI:** 10.1101/2022.05.20.492764

**Authors:** Anna Onnis, Emanuele Andreano, Chiara Cassioli, Elisa Pantano, Valentina Abbiento, Giuseppe Marotta, Rino Rappuoli, Cosima T Baldari

## Abstract

CTL-mediated killing of virally infected or malignant cells is orchestrated at a specialized intercellular junction, the immune synapse (IS). We hypothesized that SARS-CoV-2 may target IS assembly in CTLs to escape killing. We show that primary human CD8^+^ T cells strongly upregulate the expression of ACE2, the Spike protein receptor, during differentiation to CTLs. CTL pre-incubation with the Wuhan or Omicron Spike variants inhibits IS assembly and function, as shown by defective synaptic accumulation of TCRs and tyrosine phosphoproteins as well as defective centrosome and lytic granule polarisation to the IS, resulting in impaired target cell killing. These defects were reversed by anti-Spike antibodies that interfere with ACE2 binding and were reproduced by ACE2 engagement with Angiotensin-II or an anti-ACE2 antibody, but not by the ACE2 product Ang (1-7). These results highlight a new strategy of immune evasion by SARS-CoV-2 based on the Spike-dependent, ACE2-mediated targeting of the lytic IS to prevent the elimination of infected cells.

**Summary statement:** We report a new mechanism of immune evasion by SARS-CoV-2 based on direct disabling CTLs to form immune synapses through Spike protein binding to ACE2. This mechanism could contribute to the failure of the immune system to control SARS-CoV-2 infection.

## Introduction

Severe coronavirus disease 2019 (COVID-19) caused by SARS-CoV-2 infection is associated with immune dysregulation, with defective interferon type I response (Bastard et al., 2020; Zhang et al., 2020), local hyperactivation of innate immune cells and impaired adaptive immune responses, with a prominent role for T cells (Chen and John Wherry, 2020; Kalfaoglu et al., 2021). T cell defects include lymphopenia and dysfunctions that range from excessive activation and exhaustion, to defective activation, to abnormalities in differentiation (Chen and John Wherry, 2020; Kalfaoglu et al., 2021). Interestingly, CD8^+^ T cells appear preferentially dysregulated in severe COVID-19 compared to CD4^+^ T cells. A selective decrease in cytotoxic T lymphocytes (CTL) in the upper respiratory tract has been observed in patients with severe disease compared to patients with milder disease (Chua et al., 2020). Additionally, a decrease in cytokine production by CD8^+^ T cells has been reported in severe COVID-19 (Zheng et al., 2020), while a more robust CD8^+^ T cell expansion has been associated with milder disease or recovery (Liao et al., 2020; Wen et al., 2020).

As the T cell effectors responsible for the apoptotic demise of virally infected cells, CTLs are major targets for immune evasion by viral pathogens. The killing machinery of CTLs includes the lytic granules, which are lysosome-like organelles enriched in granzymes and perforin, the recently described supramolecular attack particles (Balint et al., 2020), and FasL, which is released both at the plasma membrane and in association with exosomes to trigger the Fas apoptosis pathway in target cells (Cassioli and Baldari, 2022). While the specificity of killing is dependent on cognate cell recognition by T cell antigen receptor (TCR) interaction with major histocompatibility complex-bound peptide antigen (pMHC), selectivity is ensured by the precise delivery of the cytotoxic effectors to the target cell at the immune synapse (IS), a highly specialized signaling and secretory platform that forms at the CTL interface with its target (Douanne and Griffiths, 2021; Dustin and Choudhuri, 2016). TCR engagement triggers a profound rearrangement of receptors, adhesion molecules and signaling mediators at the target cell contact, which leads to the typical bull’s eye architecture of the mature IS. A reorganization of the cytoskeleton accompanies and coordinates this process. An inward actin flow allows for the centripetal movement of TCRs and co-stimulatory receptors to the IS center, eventually forming an F-actin ring that seals a space between the CTL and its target, the synaptic cleft. Additionally, the centrosome and secretory apparatus polarise towards the synaptic membrane, with microtubules providing tracks for the transport of the cytotoxic machinery to the target cell contact and its delivery into the synaptic cleft (Blumenthal and Burkhardt, 2020; Martin-Cofreces and Sanchez-Madrid, 2018).

The differentiation of antigen-specific CD8^+^ T cells to CTLs is primed in peripheral lymphoid tissues by DCs that have been activated at the site of infection. Hence the dysregulation of the CD8^+^ T response by SARS-CoV2 is likely to result, at least in part, from defective priming. However, following their differentiation, CTLs migrate to the site of infection, where they can be exposed to high virus concentrations. The finding that the SARS-CoV-2 receptor human angiotensin-converting enzyme 2 (ACE2) (Hoffmann et al., 2020) is expressed in activated T cells and mediates binding and internalisation of SARS-CoV-2 (Welch et al., 2022) suggests that, as a result of ACE2 engagement by the Spike protein, T cells may be directly modulated by the virus.

Here we have hypothesized that, similar to lymphotropic viruses such as HIV-1 (Fackler et al., 2007), SARS-CoV-2 may target IS assembly and function as a means to escape CTL-mediated killing. We provide evidence that Spike suppresses IS assembly in CTLs, leading to impaired cytotoxicity. This effect is dependent on its interaction with ACE2 and can be reproduced by ACE2 ligands, highlighting a role for ligand-bound ACE2 as an inhibitory receptor in CTL activation and a target for immune evasion by SARS-CoV-2.

## Results and Discussion

### CTLs express the Spike receptor ACE2

While activated T cells have been reported to express the Spike receptor ACE2 (Coppo et al., 2011; Welch et al., 2022), whether this occurs in CTLs has not been investigated. We generated CTLs from CD8^+^ T cells immunopurified by negative selection from buffy coats from healthy donors and activated using beads coated with anti-CD3 and anti-CD28 mAbs in the presence of IL-2 (Fig.S1A). Under these conditions CD8^+^ T cells differentiate to functional CTLs by day 5, as assessed by granzyme B (GzmB) expression and cytotoxicity (Fig.S1B,C). Quantitative real-time RT-PCR showed that ACE2 was undetectable in freshly purified CD8^+^ T cells but was strongly upregulated in CTLs (Fig.1A).

### Spike suppresses IS formation and CTL-mediated cytotoxicity

To test the hypothesis that Spike binding to ACE2 can modulate CTL function by targeting the IS, CTLs were pre-treated for 30 min with different concentrations of the original Wuhan Spike (Spike-W) at 20°C to allow for binding but not internalization, and subsequently mixed with Raji B cells pulsed with a mix of the Staphylococcal enterotoxins A, B and E to broadly cover the TCR Vβ repertoire. This is the experimental setting typically used to image IS assembly when using polyclonal human T cells. The experiment was carried out in the absence of serum to rule out potential effects of circulating angiotensins (Ang).

The assembly of functional immune synapses was assessed by measuring the accumulation of tyrosine phosphoproteins and TCR/CD3 complexes at the T cell contact with the APC 15 min following conjugate formation, a time point when the IS has acquired its characteristic architecture. Spike was found to suppress the formation of signaling-competent immune synapses by CTLs in a dose-dependent fashion, as shown by the decreased frequency of conjugates displaying PTyr staining at the IS as well as the impairment in the synaptic accumulation of tyrosine phosphoproteins (Fig.S2A,B; Fig.1B,C; videos 1-3). Spike also suppressed TCR/CD3 accumulation at the IS (Fig.1B,D; videos 1-3), accounting for the signaling defect.

**Figure 1.**
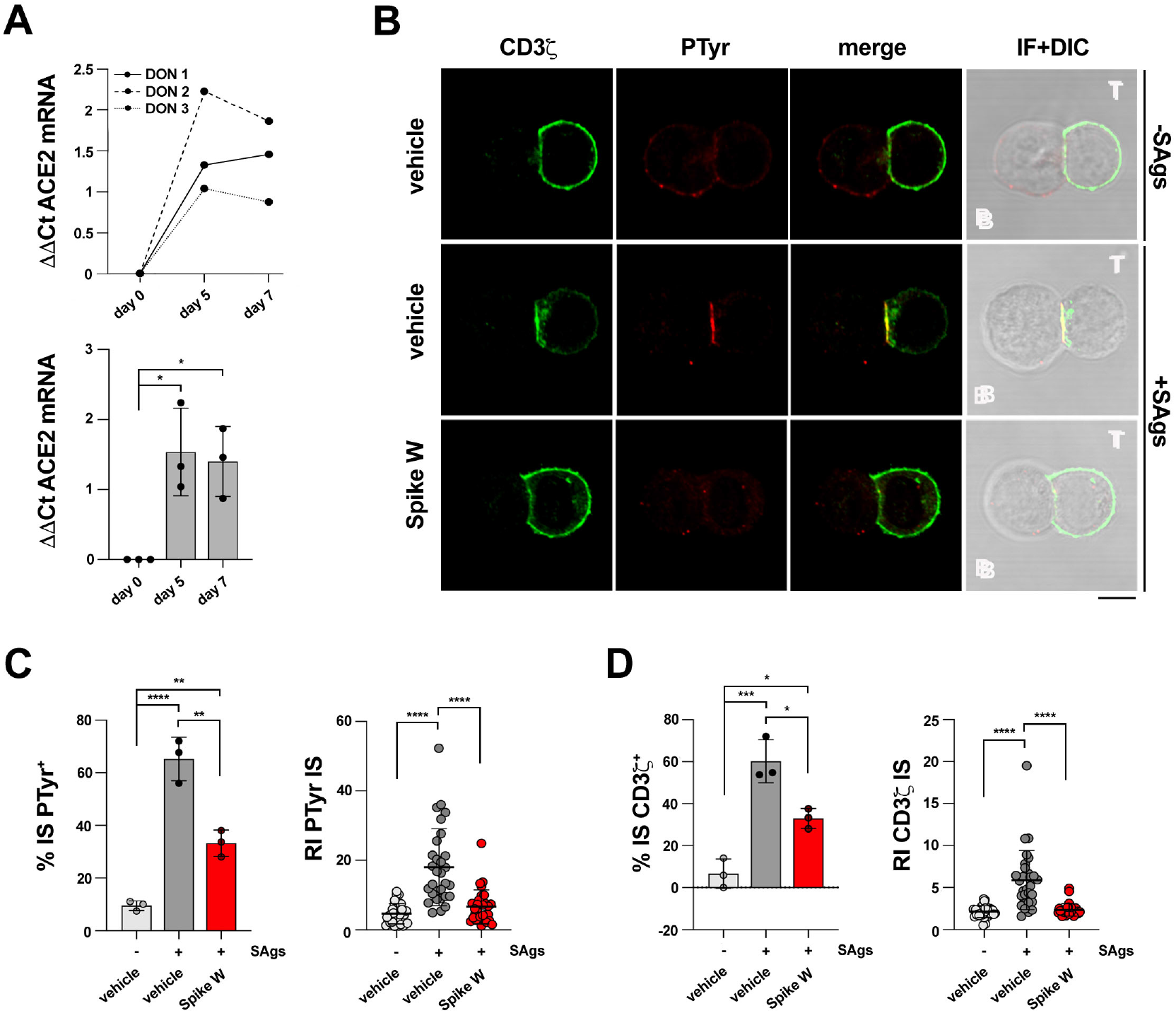
Spike suppresses TCR accumulation and phosphotyrosine signaling at the CTL IS. **A**. Quantitative RT-PCR of human ACE2 mRNA in purified CD8^+^ T cells at days 0, 5 and 7 after stimulation with anti-CD3/CD28 mAb-coated beads in the presence of IL-2 (n=3). **B-D**. Immunofluorescence analysis of CD3ζ and PTyr in CTLs (day 7) pre-treated with either vehicle (PBS) or 0.05μg/μl Spike Wuhan (Spike W; cell viability after pre-treatment 91.7±0.2%), mixed with Raji cells (APCs) either unpulsed or pulsed with a combination of SEA, SEB and SEE (SAgs), and incubated for 15 min at 37°C. Representative images (medial optical sections) of the T cell:APC conjugates are shown (**B**). **C**. *Left*, Quantification (%) of 15-min antigen-specific conjugates harboring PTyr staining at the IS (≥50 cells/sample, n=3, One-way ANOVA test). *Right*, Relative PTyr fluorescence intensity at the IS (recruitment index) (10 cells/sample, n=3, Kruskal-Wallis test). **D**. *Left*, Quantification (%) of 15-min antigen-specific conjugates harboring CD3ζ staining at the IS (≥50 cells/sample, n=3, One-way ANOVA test). *Right*, Relative CD3ζ fluorescence intensity at the IS (recruitment index) in CTLs (day 7) pre-treated with either vehicle (PBS) or 0.05μg/μl Spike Wuhan (Spike W), mixed with Raji cells (APCs) either unpulsed or pulsed with a combination of SEA, SEB and SEE (SAgs), and incubated for 15 min at 37°C (10 cells/sample, n=3, Kruskal-Wallis test). Size bar, 5 μm. \*\*\*\**p*≤0.0001; *** *p*≤0.001; \*\**p*≤0.01; \**p*≤0.05. Non-significant differences are not shown.

TCR signaling starts immediately upon T cell contact with a cognate APC and is sustained throughout the process of IS maturation. To understand whether Spike interferes with TCR signaling at an earlier stage of IS maturation we compared the outcome of CTL pretreatment with Spike-W on IS assembly in 5-min (nascent IS) versus 15-min (mature IS) conjugates. The defects in phosphotyrosine signaling and TCR/CD3 accumulation at the IS were clearly detectable in 5-min conjugates (Fig.2A,B), indicating that Spike-W impairs TCR signaling at an early step in IS formation. Consistent with this finding, the early synaptic accumulation of active ZAP-70, the protein tyrosine kinase responsible for coupling the TCR to the downstream tyrosine phosphorylation cascade (Au-Yeung et al., 2018), was compromised in Spike-W pre-treated CTLs (Fig.2C).

**Figure 2.**
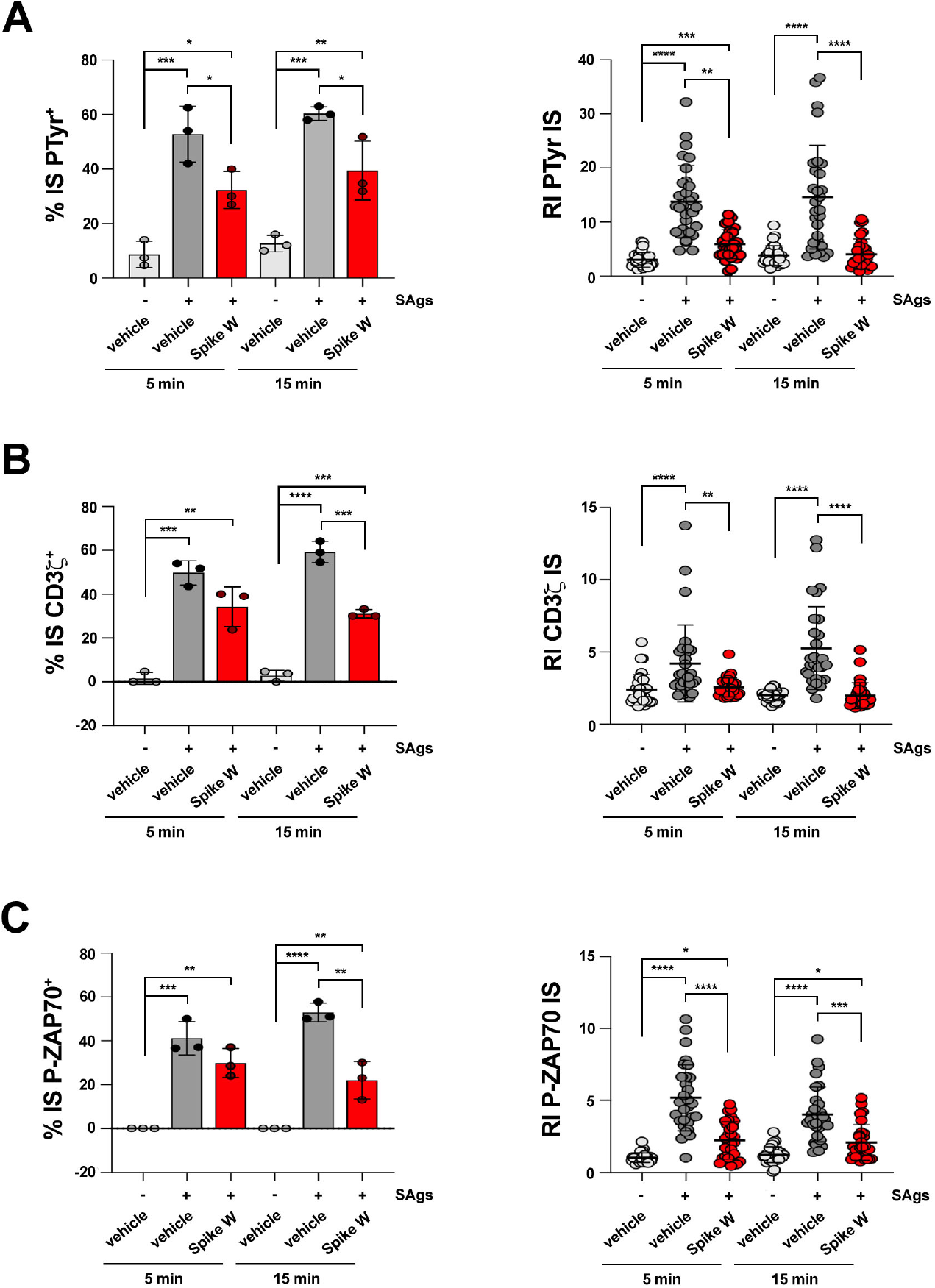
Spike inhibits centrosome and lytic granule polarisation to the CTL IS and CTL-mediated cytotoxicity. **A-C**. *Left*, Quantification (%) of 5-min and 15-min antigen-specific conjugates harboring PTyr (**A**), CD3ζ (**B**), or P-ZAP-70 (**C**) staining at the IS in CTLs (day 7) pre-treated with either vehicle (PBS) or 0.05μg/μl Spike Wuhan (Spike W), then mixed with Raji cells (APCs) either unpulsed or pulsed with a combination of SEA, SEB and SEE (SAgs), and incubated for 15 min at 37°C (≥50 cells/sample, n=3, One-way ANOVA test). *Right*, Relative PTyr (**A**), CD3ζ (**B**), or P-ZAP-70 (**C**) fluorescence intensity at the IS (recruitment index) (10 cells/sample, n=3, Kruskal-Wallis test). Data are expressed as mean±s.d. \*\*\*\**p*≤0.0001; *** *p*≤0.001; \*\**p*≤0.01; \**p*≤0.05. Non-significant differences are not shown.

Centrosome repositioning beneath the synaptic membrane is essential for the polarized delivery of the cytotoxic effectors of CTLs to the target cell (Douanne and Griffiths, 2021). Co-staining with anti-pericentrin and anti-GzmB Abs showed that centrosome polarization towards the IS was impaired in Spike-W-pretreated CTLs, as assessed by measuring the distance of the centrosome from the IS center (Fig.S2A; Fig.3A,B; videos 4-6). Additionally, the convergence of the lytic granules towards the centrosome, which is required for efficient degranulation (Daniele et al., 2011; Mentlik et al., 2010), was impaired under these conditions (Fig.S2A; Fig.3A,C; videos 4-6).

**Figure 3.**
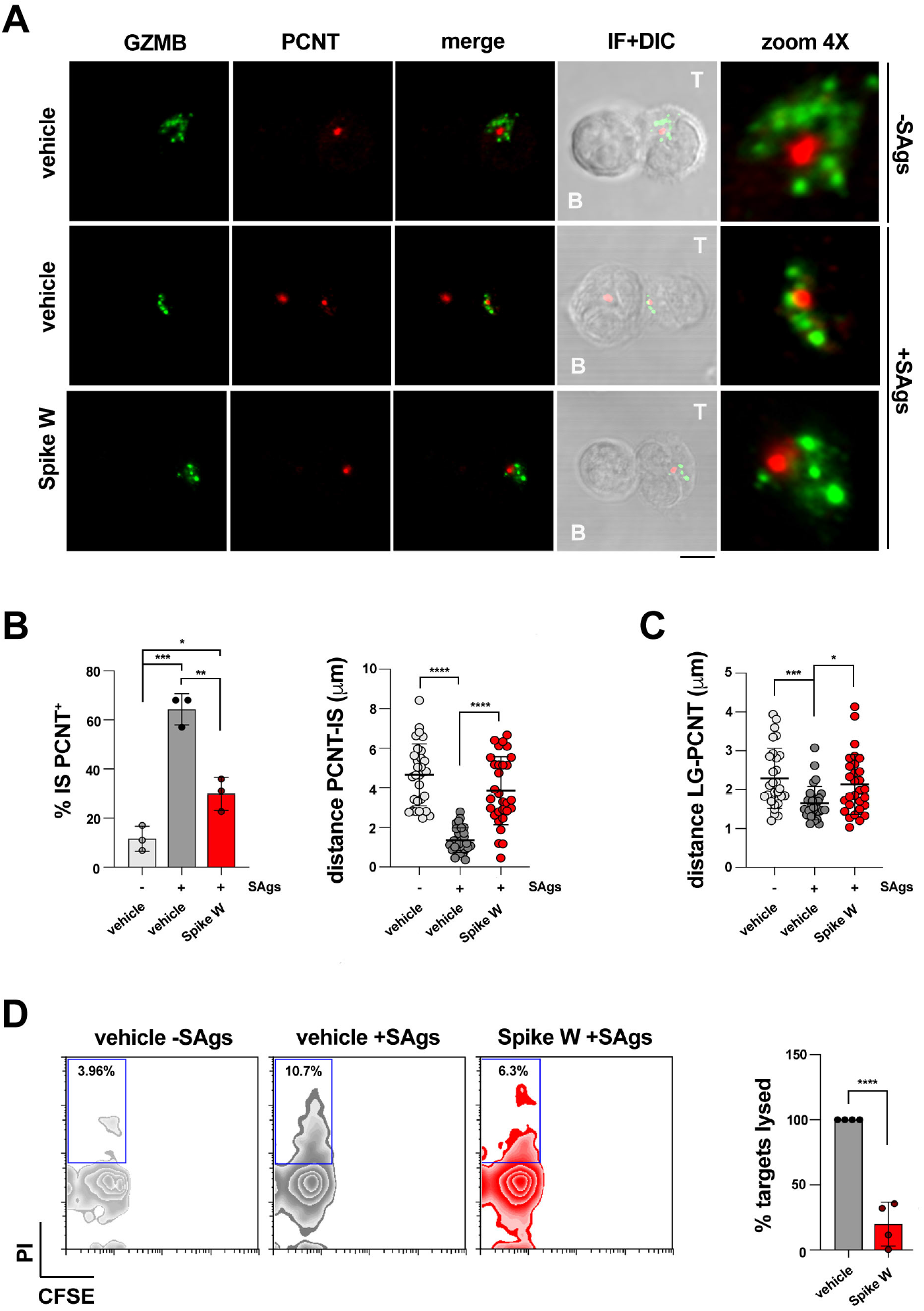
Spike inhibits centrosome and lytic granule polarisation to the CTL IS and CTL-mediated cytotoxicity. **A-C**. Immunofluorescence analysis of PCNT and GrzmB in CTLs (day 7) pre-treated with either vehicle (PBS) or 0.05μg/μl Spike Wuhan (Spike W), then mixed with Raji cells (APCs) either unpulsed or pulsed with a combination of SEA, SEB and SEE (SAgs), and incubated for 15 min at 37°C. Representative images (medial optical sections) of the conjugates are shown (**A**). **B**. *Left*, Quantification (%) of 15-min antigen-specific conjugates harboring PCNT staining at the IS (≥50 cells/sample, n=3, One-way ANOVA test). *Right*, Measurement of the distance (μm) of the centrosome ζPCNT) from the CTL-APC contact site (10 cells/sample, n=3, One-way ANOVA test). **C**. Measurement of the distance (μm) of the lytic granules ζLG, marked by GzmB) from the centrosome (PCNT) (10 cells/sample, n=3, Kruskal-Wallis test) (see figure S2 for parameters used for measurements plotted in panels B and C). **D**. Flow cytometry analysis of cytotoxicity of CTLs (day 7) pre-treated with either vehicle (PBS) or Spike Wuhan (Spike W). *Left*, Representative flow cytometric dot plot showing the percentages (%) of CFSE^-^/PI^+^ cells. *Right*, Quantitative analysis of CTL-mediated cytotoxicity of Raji B cells. The histograms show the percentage (%) of target cells lysed (n=4, unpaired two-tailed Student’s *t*-test). Values for the treatment with vehicle were set as 100%. Size bar, 5 μm. Data are expressed as mean±SD. \*\*\*\**p*≤0.0001; *** *p*≤0.001; \*\**p*≤0.01; \**p*≤0.05. Non-significant differences are not shown.

To test the outcome of this defect on CTL-mediated killing, we carried out cytotoxicity assays by incubating Spike-W-pretreated, CFSE-labelled CTLs with SAg-pulsed Raji cells for 4 h, followed by staining with propidium iodide and flow cytometric analysis, gating on the CFSE-negative cell population. Consistent with the defect in lytic granule transport to the IS, Spike-W was found to suppress the ability of CTLs to kill target cells (Fig.3D). Hence Spike-W prevents CTLs from forming functional immune synapses, thereby compromising their killing potential.

### The inhibitory effects of Spike-W on IS formation are reproduced by the Omicron BA.1 and BA.2 variants and are reversed by antibodies that prevent Spike binding to ACE2

To achieve insights into the mechanism underlying the suppressive activity of Spike on CTL IS assembly and function, we extended the analysis to the recent Spike variants Omicron BA.1 and BA.2. These variants recapitulated the effects of Spike-W on IS assembly, as assessed by measuring the synaptic accumulation of tyrosine phosphoproteins and TCR/CD3 complexes (Fig.4A,B), as well as the polarization of the centrosome toward the IS and the convergence of lytic granules around the centrosome (Fig.4C,D). Since the mutations carried by the Omicron Spike variants do not compromise ACE2 binding (Wrapp et al., 2020; Yin et al., 2022), this result suggests that the effects of Spike on CTLs are mediated by ACE2. Consistent with this notion, pre-treatment of CTLs with Spike-W or Spike Omicron BA.1 in the presence of specific neutralizing mAbs that prevent Spike binding to ACE2 (Andreano et al., 2021a; Andreano et al., 2021b; Torres et al., 2022) reverted the suppressive effects of the Spike variants on IS assembly by CTLs (Fig.4A-D). Additionally, no effects of Spike on IS assembly were detectable in freshly purified CD8^+^ T cells (Fig.S2C-E), which do not express ACE2 (Fig.1A).

**Figure 4.**
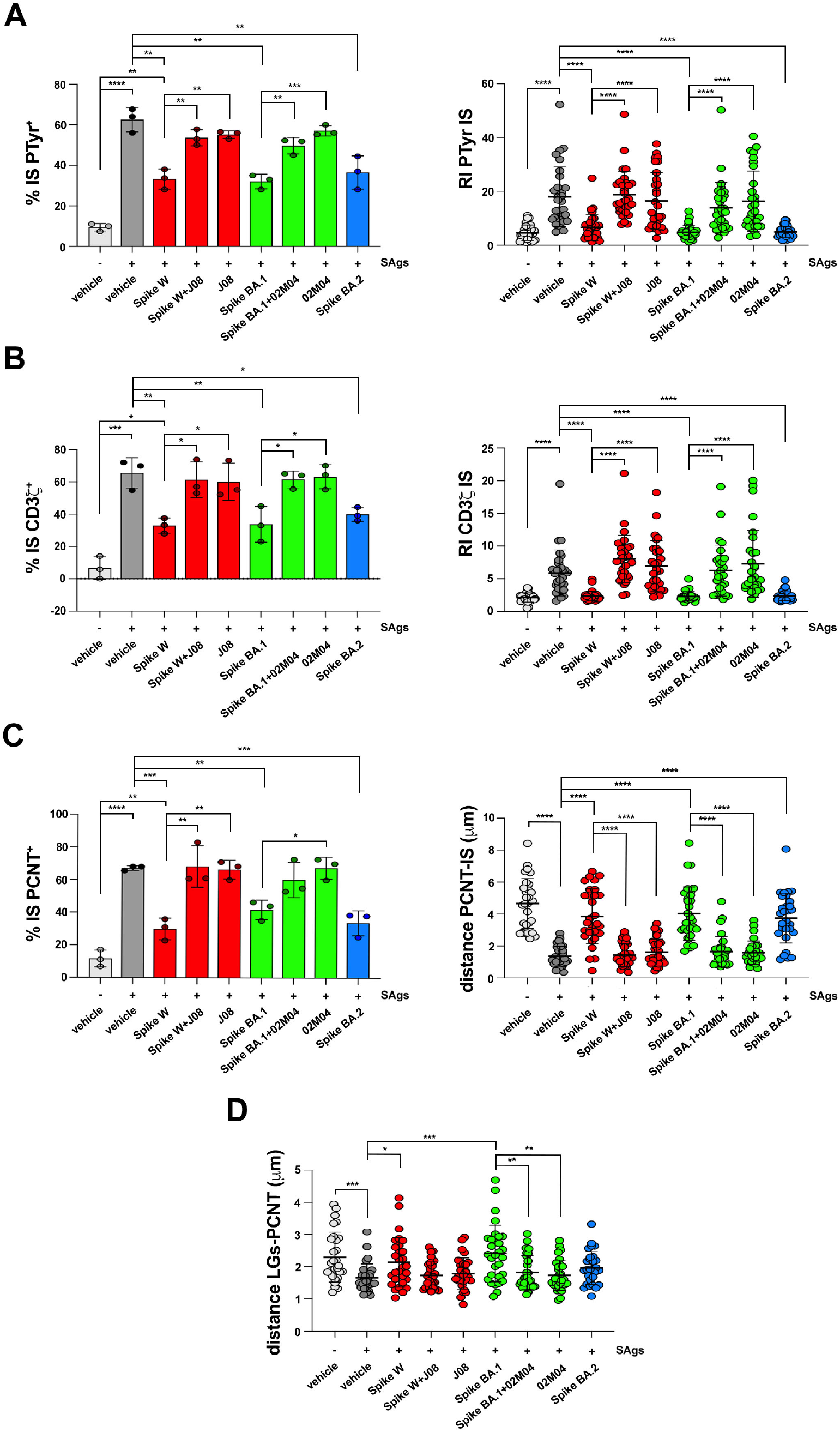
Inhibition of CTL IS assembly by Spike-W or the Omicron BA.1 and BA.2 variants can be reversed by mAbs that interfere with ACE2 binding. **A**,**B**. *Left*, Quantification (%) of 15-min antigen-specific conjugates harboring PTyr (**A**) or CD3ζ (**B**) staining at the IS in CTLs (day 7) pre-treated with either vehicle (PBS) or 0.05μg/μl Spike Wuhan (Spike W), Spike Omicron BA.1 (Spike BA.1), alone or in the presence of the respective neutralizing mAb (Spike W+J08; Spike BA.1+02M04); or Spike Omicron BA.2 (Spike BA.2). Samples pre-treated with only the neutralizing mAbs (J08 and 02M04) were also included in the analysis. CTLs were mixed with Raji cells (APCs), either unpulsed or pulsed with a combination of SEA, SEB and SEE (SAgs), and incubated for 15 min at 37°C (≥50 cells/sample, n=3, unpaired two-tailed Student’s *t*-test). *Right*, Relative PTyr (**A**) or CD3ζ (**B**) fluorescence intensity at the IS (recruitment index) (10 cells/sample, n=3, Kruskal-Wallis test). **C**. *Left*, Quantification (%) of 15-min antigen-specific conjugates formed as in panel A harboring PCTN1 staining at the IS (≥50 cells/sample, n=3, One-way ANOVA test). *Right*, Measurement of the distance (μm) of the centrosome ζPCNT) from the T cell-APC contact site (10 cells/sample, n=3, Kruskal-Wallis test). **D**. Measurement of the distance (μm) of the lytic granules ζLG, marked by GzmB) from the centrosome (PCNT) in 15-min antigen-specific conjugates formed as in panel A (10 cells/sample, n=3, Kruskal-Wallis test). \*\*\*\**p*≤0.0001; *** *p*≤0.001; \*\**p*≤0.01; \**p*≤0.05. Non-significant differences are not shown.

### The inhibitory effects of Spike on IS formation are mediated by ACE2

ACE2 is a type I transmembrane zinc metalloprotease that plays a key role in the renin-angiotensin system (RAS). The extracellular carboxypeptidase domain of ACE2 catalyses the conversion of Angiotensin (Ang)-I to Ang (1-9), and Ang-II and Ang (1-9) to the bioactive peptide Ang (1-7) which activates the receptor MAS1 on target cells (Samavati and Uhal, 2020). However, ACE2 also acts as a receptor that modulates intracellular processes in a peptidase-independent manner through distinct contributions of its extracellular, transmembrane and intracellular domains, including integrin transactivation, regulation of aminoacid transporters and endocytosis-dependent infection by SARS-CoV-2 (Danilczyk et al., 2006; Hoffmann et al., 2020; Inoue et al., 2007; Simons et al., 2021). The fact that mAbs that prevent Spike interaction with ACE2 reverse the effects of Spike supports the notion that the inhibitory effects on IS formation are mediated by ACE2 signaling. To test this hypothesis we assessed the outcome of ACE2 engagement on IS assembly in CTLs, using as ligand an anti-ACE2 polyclonal Ab raised against the extracellular domain that blocks SARS-CoV-2 entry in target cells by competing with Spike for ACE2 binding (Hoffmann et al., 2020). CTL pre-treatment with the anti-ACE2 Ab recapitulated the suppressive effects of Spike on IS assembly, including the synaptic accumulation of tyrosine phosphoproteins and TCR/CD3 complexes (Fig.5A,B), centrosome polarization toward the IS (Fig.5C) and lytic granule convergence around the centrosome (Fig.5D). Consistently, CTL-mediated killing was suppressed under these conditions (Fig.5E). Similar to Spike, the anti-ACE2 Ab suppressed the formation of signaling-competent immune synapses in CTLs at an early step of IS maturation, as assessed by analyzing immune synapses formed in 5-min versus 15-min CTL-target cell conjugates (Fig.S3A-C). No effect on IS assembly was observed when freshly purified CD8^+^ T, which do not express ACE2 (Fig.1A), were pre-treated with the anti-ACE2 Ab (Fig.S3D-F). Of note, while AngII suppressed the assembly of functional immune synapses in CTLs, similar to the anti-ACE2 Ab, no effect was observed when cells were treated with Ang (1-7) (Fig.5A-D), ruling out the catalytic activity of ACE2 in this function as well as indirect effects mediated by the Ang (1-7) receptor MAS1. In this context, although we found MAS1 to be expressed in CD8^+^ T cell, its expression decreased during their differentiation to CTLs (Fig.S3G). Taken together, these results support the notion that, following ligand binding, ACE2 delivers inhibitory signals to suppress IS assembly.

**Figure 5.**
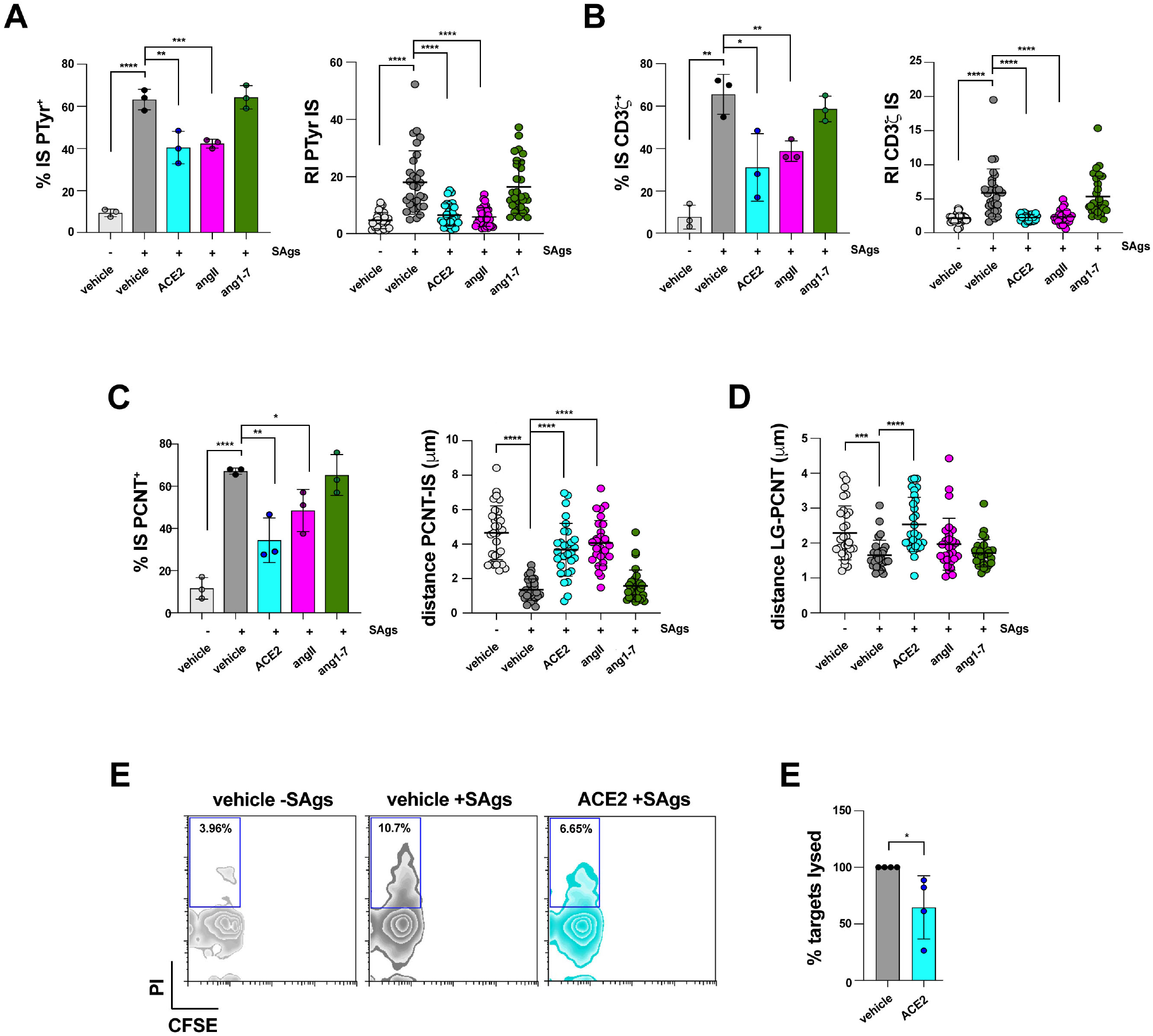
The inhibitory effects of Spike on IS formation are mediated by ACE2. **A**,**B**. *Left*, Quantification (%) of 15-min conjugates harboring PTyr (**A**) or CD3ζ (**B**) staining at the IS. CTLs (day 7) were conjugated with Raji cells (APCs) in the absence or presence of SAgs and either an anti-ACE2 Ab (cell viability after pre-treatment 93.7±0.8%), or angiotensin II (AngII), or the peptide angiotensin 1-7 (Ang 1-7) (≥50 cells/sample, n=3, One-way ANOVA test). *Right*, Relative PTyr (**A**) or CD3ζ (**B**) fluorescence intensity at the IS (recruitment index) (10 cells/sample, n=3, Kruskal-Wallis test). **C**. *Left*, Quantification (%) of 15-min conjugates formed as in panel A harboring PCTN staining at the IS (≥50 cells/sample, n=3, One-way ANOVA test). *Right*, Measurement of the distance (μm) of the centrosome ζPCNT) from the T cell-APC contact site in CTL-APC conjugates formed as in panel A (10 cells/sample, n=3, Kruskal-Wallis test). **D**. Measurement of the distance (μm) of the lytic granules ζLG, marked by GzmB) from the centrosome (PCNT) in 15-min CTL-APC conjugates formed as in panel A (10 cells/sample, n=3, Kruskal-Wallis test). **E**. *Left*, Flow cytometry analysis of cytotoxicity of CTLs (day 7) pre-treated with either vehicle (PBS) or 2 μg/ml anti-ACE2 Ab (ACE2). *Left*, Representative flow cytometric dot plot showing the percentages (%) of CFSE^-^/PI^+^ cells. *Right*, Quantitative analysis of CTL-mediated cytotoxicity of Raji B cells. The histograms show the percentage (%) of target cells lysed (n=4, unpaired two-tailed Student’s *t*-test). Values for the treatment with vehicle were set as 100%. The data are expressed as mean±SD. \*\*\*\**p*≤0.0001; *** *p*≤0.001; \*\**p*≤0.01; \**p*≤0.05. Non-significant differences are not shown.

Collectively, our data provide novel insight into the mechanisms of immune evasion by SARS-CoV-2, involving the Spike-ACE2 axis, as a means for infected cells to avoid killing. The precise mechanism underlying the inhibitory effects of Spike on IS assembly remains to be elucidated. Our data show that SARS-CoV-2 Spike protein binding to ACE2 interferes with the assembly of a functional IS in CTLs by inhibiting the process at an early step. Of note, the Omicron variants suppress IS assembly to a similar extent as Spike-W, despite a higher binding affinity for ACE2 (Wrapp et al., 2020; Yin et al., 2022), suggesting that Spike-W binding to ACE2 is sufficient to fully unleash the inhibitory potential of ACE2 engagement on this process. Potential targets of the ACE2-Spike axis are integrins such as LFA-1, which is essential for the generation of well-structured immune synapses characterised by the tight accumulation of TCR/CD3 complexes at the center of the contact area with the APC (Cassioli et al., 2021a). ACE2 has been reported to interact with β1 integrin, a fibronectin receptor, inducing its conversion to the open, high-affinity conformation and activation of downstream signaling mediators such as FAK (Clarke et al., 2012; Lin et al., 2004). Whether ACE2 modulates integrin signaling in T cells is an interesting hypothesis that warrants testing.

It is noteworthy that, close to the ACE2 binding site, Spike displays a conserved RGD sequence, the consensus integrin-binding motif (Sigrist et al., 2013). Experimental evidence supports the notion that β1 and αv integrins may act as alternative Spike receptors or co-receptors together with ACE2 (Norris et al., 2022; Park et al., 2021). Mechanistically, Spike has been shown to modulate β1 integrin signaling in the kidney cell line Vero E6 for productive infection by activating inside-out signaling mediated by Gα13 and the integrin adaptor talin (Simons et al., 2021). Additionally, the RGD sequence within the ACE2-binding domain of Spike has been recently identified as an αv-selective integrin agonist in primary human airway epithelial cells, where it triggers cell spreading, formation of focal adhesions and actin reorganisation through the activation of the kinases FAK and Src, and the adaptor paxillin (Norris et al., 2022). However, the fact that integrin activation is a key activating event required for the onset of IS assembly (Springer and Dustin, 2012), while Spike suppresses this process, strongly suggests that the inhibitory effects of Spike on the CTL IS are unlikely to be mediated by integrins. Additionally, our data show that the Spike Omicron BA.2 variant, which carries a non-conservative D to N substitution at position 405 of the RGD integrin binding motif mapping at positions 403-405 (RGN), is as effective as Spike-W and Spike Omicron BA-1 at suppressing IS assembly. Conversely, the ability of mAbs that prevent Spike binding to ACE2, together with the lack of any effect of Spike on IS assembly in freshly purified CD8^+^ T cells that do not express ACE2, highlights the key role of ACE2 in the Spike-dependent suppression of IS assembly and CTL-mediated killing.

The finding that ACE2 engagement by the ACE2 ligand Ang-II, or by an anti-ACE2 Ab specific for the extracellular domain, elicits effects that recapitulate those observed for Spike on the CTL IS further supports the notion that, upon engagement by ligand, ACE2 acts as a receptor that negatively regulates the TCR-derived signals driving CTL IS assembly. While the role of ACE2 as an ectoenzyme that participates in the RAS has been extensively characterized, the role of its transmembrane and intracellular domains, that together display 48% identity with collectrin (Zhang et al., 2001), are only beginning to be explored. Both domains have been implicated in SARS-CoV-2 infection through the endocytic uptake of the virus following Spike-mediated binding to the extracellular domain of ACE2 (Hoffmann et al., 2020; Inoue et al., 2007). Additionally, the collectrin homology domain has been shown to act as a chaperone to stabilise transporters of large neutral aminoacids at the surface of intestinal epithelial cells to regulate aminoacid absorption (Camargo et al., 2009; Kowalczuk et al., 2008). The intracellular domain also regulates the TACE/ADAM17-mediated shedding of the extracellular domain to its catalytically active soluble form by interacting with calmodulin (Lambert et al., 2008). Whether these peptidase-independent functions may be relevant to CTL IS assembly and target cell killing remains to be investigated.

It is noteworthy that a functional RAS has been documented in T cells (Coppo et al., 2011). Among the RAS components, the Ang-II receptor AT1 has been reported to have immunomodulatory effects on T cells, limiting harmful CD8^+^ T cell responses during blood-stage malaria and promoting a protective CD4^+^ T cell response in the kidney in the setting of hypertension (Silva-Filho et al., 2016; Silva-Filho et al., 2017; Zhang et al., 2012). This suggests that ACE2 may participate in T cell immunity through both its enzyme-independent and its enzyme-dependent activities.

Other viruses have co-opted the T cell IS as a target to escape immunosurveillance. One of the most striking examples is HIV-1, which exploits the virulence factor Nef to subvert the process of vesicular trafficking required for the coordinated transport of key signaling components to the IS. The resulting IS abnormalities and signaling defects eventually lead to a nonproductive T cell response against the virus (Fackler et al., 2007). Importantly, the IS acts a signaling and cell-cell communication platform also for other immune cells, including helper T cells and B cells (Batista et al., 2001; Chemin et al., 2012; Papa and Vinuesa, 2018). The data reported here highlight the intriguing possibility that SARS-CoV-2 may co-opt IS targeting as a wide-ranging immunomodulatory strategy, beyond suppression of CTL function, to evade the adaptive immune response of the host.

## Materials and Methods

### Cells and antibodies

Primary human CD8^+^ T cells were isolated from buffy coats of healthy donors and purified (>95%) by negative selection using the RosetteSep™ Human CD8^+^ T Cell Enrichment Cocktail (StemCell Technologies, #15063), according to the manufacturer’s instructions. CD8^+^ T cells were cultured immediately following isolation at 37°C, 5% CO_2_, in RPMI-HEPES medium (Merck, #R7388) supplemented with 10% iron-enriched bovine calf serum (BCS; GE Healthcare HyClone, #SH30072.03) and 1% MEM Non-essential amino acids (MEM NEAA) (#11140050) at the density of 1×10^6^/ml. The Burkitt Lymphoma derived B cell line Raji was grown at 37°C, 5% CO_2_, in RPMI-1640 Medium (Merck, #R8758) supplemented with 7.5% BCS.

Primary commercial antibodies used in this work were from SantaCruz (CD3ζ, clone 6B10.2, #sc-1239), Cell Signaling (P-ZAP-70 Y319/ Syk Y352, #2701S; PTyr, #8954S), Abcam (Pericentrin, #4448) and BD Biosciences (Granzyme B, #560211). Alexa Fluor 488-and 555-labeled secondary antibodies were from ThermoFisher Scientific (anti-mouse 488, #A11001; anti-rabbit 555, #A21428).

### SARS-CoV-2 Spike protein expression and purification

Plasmids encoding the Wuhan Spike (Spike-W) and Spike Omicron BA.1 and BA.2, were transiently transfected in Expi293F™ cells and ExpiCHO-S cells (Thermo Fisher) respectively. Cells were grown for six days at 37 °C with 8% CO_2_ shaking 125 rpm according to the manufacturer’s protocol (Thermo Fisher). Cell cultures for Spike-W and Spike Omicron BA.1 and BA.2 were harvested six days after transfection. Collected supernatants were clarified by centrifugation (1,200 x g, 30 min, 4°C) followed by filtration through a 0.45 μm filter. Chromatography purification was conducted at room temperature using ÄKTA go purifier system from GE Healthcare Life Sciences. Filtrated culture supernatants were purified with a 5 ml HisTrap FF Crude column (GE Healthcare Life Sciences) previously equilibrated in Buffer A (20 mM NaH_2_PO_4_, 500 mM NaCl + 30 mM Imidazol pH 7.4). Spike proteins were eluted from the column with 5 column volumes of 60% Buffer B (20 mM NaH_2_PO_4_, 500 mM NaCl + 500 mM Imidazol pH 7.4). Eluted fractions were pooled and dialyzed with PBS buffer pH 7.4 using Slide-A-Lyzer™ Dialysis Cassette 10K MWCO (Thermo Scientific) overnight at 4°C. The final Spike protein concentration was determined by measuring the absorbance at 562 nm using Pierce™ BCA Protein Assay Kit (Thermo Scientific). Final proteins were dispensed in aliquots of 0.5 ml each and stored at -80°C prior to use.

### Expression and purification of human monoclonal antibodies

Expi293F™ cells (Thermo Fisher) were transfected with plasmids carrying J08 and 02M04 antibody heavy and the light chains with a 1:2 ratio using ExpiFectamineTM293 Transfection Kit (Thermo Fisher Scientific) as recommended by the manufacturer. Three and six days after transfection, cell cultures were harvested and clarified by centrifugation (1,100 g for 10 min at RT). Supernatants were recovered and filtered with 0.45 μm filter to remove particulate material. Purification process was conducted at room temperature using the ÄKTA Go purification system (GE Healthcare Life Sciences) through a 1 ml HiTrap Protein G HP column (GE Healthcare Life Sciences) previously equilibrated with Loading Buffer (0.02 M NaH2PO4 pH 7). Each monoclonal antibody was eluted from the column in 1 ml fractions of Elution Buffer (0.1 M glycine-HCl, pH 2.7) and collected in vials pre-dispensed with 100 μl of Neutralization Buffer (Tris-HCl pH 9.0). Protein-containing fractions were pooled and dialyzed in PBS 1x Buffer pH 7.4 with a 1:200 ratio using Slide-A-Lyzer™ Dialysis Cassettes, 10K MWCO, 3 mL (Thermo Fisher Scientific) at 4°C overnight. The final antibody concentration was determined by measuring the absorbance at 562 nm using Pierce™ BCA Protein Assay Kit (Thermo Scientific). Purified antibodies were stored at -80°C prior to use.

### CTL differentiation

CD8^+^ T cells (1×10^6^/ml) were stimulated and expanded in RPMI-HEPES supplemented with 10% BCS, 1% MEM NEAA and 50 Units/mL recombinant human IL-2 (Myltenyi; #130-097-745) with Dynabeads™ Human T-activator CD3/CD28 (Gibco, #11132D) the same day of the purification (day 0, also referred as resting CD8^+^ T cells). 48 h after activation (day 2) the beads were removed and the cells expanded in RPMI-HEPES supplemented with 10% BCS, 1% MEM NEAA and 50 Units/mL recombinant human IL-2 for further three days (day 5). Five days post-purification, cells were further expanded for two days and collected at the day 7.

### CTL pre-treatments

To allow for Spike binding on the cell surface, CD8^+^ T cells (0.5×10^6^) were incubated for 30 min at 20°C in 50 μl RPMI-HEPES in the absence of BCS with 2.5 μg Spike Wuhan/BA.1/BA.2 or PBS vehicle control (concentration of Spike treatment chosen based on a dose-response analysis; see Figure S2B). Spike Wuhan and the variant Omicron BA.1 were also incubated in combination with the respective neutralizing mAbs (J08 and 02M04, respectively) at a 1:3 protein:antibody ratio. Alternatively, CD8^+^ T cells (0.5×10^6^) were incubated for 30 min at 20°C in 50 μl RPMI-HEPES in the absence of BCS with 2 μg/ml anti-ACE2 antibody (R&D Systems, #AF933) (Hoffmann et al., 2020). Alternatively, cells were incubated with the ACE2 substrate/ligand Angiotensin II (Merck, #A9525; 1 μM) or the ACE2 product Ang (1-7) (Sigma, #A9202; 1 μM) in the same culture conditions described above.

For cytotoxicity assays, cells were pre-treated in the same conditions in serum-free AIM V medium (Gibc0, #12055-091). None of the treatments affected cell viability at the concentrations and times chosen for the analyses, as assessed by Trypan blue (Sigma-Aldrich, T8154) exclusion (see legend to figures 1 and 5).

### Conjugate formation, immunofluorescence microscopy and analysis

Conjugates between CD8^+^ T cells and superantigens-pulsed Raji B cells were carried out as previously described (Cassioli et al., 2021b). Raji B cells (0.08×10^6^), used as APCs, were loaded with 10 μg/ml Staphylococcal superantigens A (SEA; Toxin Technologies, #AT101), B (SEB; Toxin Technologies, #BT202) and E (SEE; Toxin Technologies, #ET404) for 2 h and labelled with 10 μM Cell Tracker Blue for the last 15 min of the incubation with the superantigens (+ SAgs). The mix of SAgs was used to cover a substantial proportion of the TCR Vβ repertoire. Antigen-independent conjugates of CD8^+^ T cells with Raji B cells formed in the absence of SAgs were used as negative controls (-SAgs). Raji B cells, pulsed or not with SAgs, were washed twice, mixed with CD8^+^ T cells (0.12 ×10^6^) pre-treated as described above and incubated for 15 min at 37°C. To analyze early TCR signaling events a shorter time of incubation (5 min) was included. Samples for immunofluorescence analysis were seeded onto poly-L-lysine (Merck, #P1274)-coated slides (ThermoFisher Scientific, #X2XER208B) and fixed and permeabilized for 10 min in methanol at -20°C (for CD3ζ and PTyr staining) or fixed for 15 min with 4% paraformaldehyde/PBS at room temperature (for PCNT, P-ZAP70 and GzmB staining). Following fixation, samples were washed in PBS for 5 min. PFA-fixed cells were permeabilized with 0.1% Triton, 1% BSA PBS. Following permeabilization, samples were washed in PBS for 5 min. Cells were stained with primary antibodies ([1:30] CD3ζ; [1:50] P-ZAP70; [1:100] PTyr; [1:200] PCNT; [1:50] GzmB; see above catalogue #) overnight at 4°C. After washing with PBS, samples were incubated for 45 min at room temperature with Alexa Fluor 488-and 555-labeled secondary antibodies and mounted with 90% glycerol/PBS. Confocal microscopy was carried out on a Zeiss LSM700 (Carl Zeiss, Jena, Germany) microscope using a 63x/1.40 oil immersion objective. Images were acquired with pinholes opened to obtain 0.8 μm-tick sections. Detectors were set to detect an optimal signal below the saturation limits. Images were processed with Zen 2009 image software (Carl Zeiss, Jena, Germany). Immunofluorescence analyses were carried out on medial optical sections of CD8^+^ T cell:APC conjugates using ImageJ (version 1.53a). Relative distances (μm) of the centrosome (marked by PCNT) from the center of the contact site with the APC, and of the lytic granules (marked by GzmB) from the centrosome, were measured using ImageJ (FigS2A). Scoring of conjugates for accumulation of CD3ζ or P-Tyr at the IS, or for centrosome (PCNT) juxtaposition to the IS membrane, was based on the presence of the respective staining solely at the T cell-APC contact site and was expressed as percentage of conjugates with synaptic staining *versus* the total number of conjugates analyzed. Recruitment index was calculated for each marker as the ratio of CD3ζ or p-Tyr fluorescence intensity at the synaptic area, which is manually defined at the T cell-APC contact site, compared to the remaining T-cell area, using ImageJ. Z-stacks were acquired on a spinning disk confocal and super-resolution microscope (CSU-W1-SoRA Nikon), with 60x oil objectives (numerical aperture 1.49) using a Photometrics BSI (Nikon). 3D Deconvolution (Richardson-Lucy method, 10 iterations) was applied to high-resolution imaging to improve contrast of video 2 and 3. 3D reconstructions were obtained using Fiji (version 2.1.0).

### RNA purification and RT-PCR

RNA was extracted from CTLs (days 0, 5 and 7) using the RNeasy Plus Mini Kit (Qiagen. #74136), reverse transcribed to first-strand cDNAs using iScript™ cDNA Synthesis Kit (Bio-Rad, #1708891) and analyzed by Real-time quantitative PCR (RT-qPCR) as previously described (Onnis et al., 2015). Primer sequences for ACE2 amplified a region of 133 bp (Forward: TCCATTGGTCTTCTGTCACCCGC; Reverse: AGACCATCCACCTCCACTTCTC). Primer sequences for MAS1 amplified a region of 121 bp (Forward: GCCCTGAGGAGACCGTAG; Reverse: ACAACAGCGGTTCTTGCTC. Primer sequences for GrzmB amplified a region of 121 bp (Forward: TCAAAGAACAGGAGCCGACC; Reverse: TTGGCCTTTCTCTCCAGCTG). Primer sequences for 18S amplified a region of 64 bp: Forward: CGCCGCTAGAGGTGAAATT; Reverse: CTTGGCAAATGCTTTCGC. Differences in gene expression were calculated using the ΔΔCt method and normalized to 18S.

### Cytotoxicity assay

For the cytotoxicity assay, Raji B cells were loaded with 2 μg/ml Staphylococcal superantigens A, B and E (+ SAgs) for 2 h in serum-free AIM V medium (Gibco, #12055-091). Unpulsed Raji B cells were used as negative control (-SAgs). To measure cytotoxicity in *in vitro*-differentiated CTLs the calcein-based assay was used (Kummerow et al., 2014). Calcein-AM (Invitrogen, #C1430)-loaded B cells (0.050×10^6^) were incubated with CD8^+^ T cells at different ratios (1:1, 1:5, 1:10 and 1:20) in 250μl

AIMV medium for 4 h at 37°C. Reading was performed at 485 nm excitation wavelength and 528 nm emission wavelength in the bottom reading mode by a Synergy HTX multi-mode plate reader (BioTek). Cytotoxicity (% target cell lysis) was calculated as follows: (F_live_ − γ × F_exp_)/(F_live_ − Flyse) × 100, where F_live_ is the fluorescence of only target cell controls, F_exp_ are the CTLs with target cells, and F_lyse_ is the maximal target lysis. The γ value was measured at time zero: γ = F_live_ (0)/F_exp_ (0).

To analyse the effects of CTL pre-treatments on cytotoxicity, B cells (0.025×10^6^) were incubated with carboxyfluorescein diacetate succinimidyl ester (CFSE,ThermoFisher, #C34554)-loaded CD8^+^ T cells (0.225 ×10^6^), pre-treated as described above, at a 1:9 ratio in 50 μl AIMV medium for 4 h at 37°C. Cells were then diluted to 200 μl with cold PBS and acquired using a GUAVA flow cytometer (Merck Millipore). Propidium Iodide (PI,Sigma, #537059) was added before of each acquisition. Cytotoxicity (% target cell lysis) was calculated as follows: (% CFSE^−^ PI^+^ dead cells − % CFSE^−^ PI^+^ dead cells in control sample)×100/(100 − % CFSE^−^ PI^+^ dead cells in control sample).

### Statistics and reproducibility

Each experiment was performed at least 3 independent times. The exact number of repeats and the number of cells analyzed is specified in the figure legends. Statistical analyses were performed with Prism software (GraphPad Software). Pairwise or multiple comparisons of values with normal distribution were carried out with Student’s *t*-test (paired or unpaired), one-sample *t*-test (theoretical mean = 1) and One-way ANOVA test or One-sample *t*-test (theoretical mean =100), whereas values without Gaussian distribution were analysed with Mann-Whitney test or Kruskal-Wallis test. Statistical significance was defined as: \*\*\*\**p*≤0.0001; *** *p*≤0.001; \*\**p*≤0.01; \**p*≤0.05; n.s., not significant.

### Online Supplemental material

Included in the supplemental material are three supplementary figures showing the workflow for CTL generation (Fig.S1), a comparison of the effects of Spike-W on IS signaling in freshly purified and day-7 CD8^+^ T cells (Fig.S2), and the effects of anti-ACE2 mAb on the nascent and mature IS in CTLs (Fig.S3). Additionally, the supplemental material includes six videos. Videos 1-3 show the 3-D reconstruction of a representative CTL-APC conjugate formed with either SAg-unpulsed (video 1) or SAg-pulsed (videos 2 and 3) APCs and co-stained for PTyr and CD3ζ. CTLs used in SAg-specific conjugates were pre-treated with either vehicle (video 2) or Spike-W (video 3). Videos 4-6 show the 3-D reconstruction of representative CTL-APC conjugate formed with either SAg-unpulsed (video 4) or SAg-pulsed (videos 5 and 6) APCs and co-stained for PNCT and GzmB. CTLs used in SAg-specific conjugates were pre-treated with either vehicle (video 5) or Spike-W (video 6).

## Author contributions

AO, EA, CC, EP and VA did the experiments; AO and CC analysed the data and prepared the figures; AO, EA, RR and CTB wrote the paper; GM provided essential reagents.

## Acknowledgements

This work was funded by ERC Synergy grant 951329 (ATTACK) and AIRC grant IG-2017-20148 to CTB, and ERC Advanced grant 787552 to RR (vAMRes). This work was supported by a fundraising activity promoted by Unicoop Firenze, Coop Alleanza 3.0, Unicoop Tirreno, Coop Centro Italia, Coop Reno e Coop Amiatina. The graphical abstract was created using BioRender.com.

## List of non-standard abbreviations

ACE2, angiotensin-converting enzyme; Ang, angiotensin; CFSE, carboxyfluorescein succinimidyl ester; GzmB, Granzyme B; IS, immune synapse; MEM NEAA, MEM non-essential aminocids; PCNT, pericentrin; RAS, renin-angiotensin system; SEA, SEB, SEE, Enterococcal Superantigen A, B, E; SAgs, Superantigens.

