## Supplemental material for "SARS-CoV-2 Spike protein suppresses CTL-mediated killing by inhibiting immune synapse assembly"

**Figure S1. Workflow for CTL generation. A.** Workflow for CTL generation from $CD8^+$ T cells purified from buffy coats of healthy donors. **B**. Quantitative RT-PCR of human GzmB mRNA in purified $CD8^+$ T cells at days 0, 5 and 7 after stimulation with anti-CD3/CD28 mAb-coated beads in the presence of IL-2 (n=5, One-way ANOVA test). **C**. Fluorimetric analysis of cytotoxicity of CTLs (day 7) generated as depicted in panel A, using the calcein release assay. CTLs were co-cultured with SAg-pulsed Raji B cells at different CTL/target cell ratios for 4 h. The histogram shows the percentage (%) of target cells lysed (n=3, One-sample *t* test and Wilconox test). Values for the CTL/target cell ratio 1:20 were set as 100%. Data are expressed as mean±SD. ****$p \leq 0.0001$; *** $p \leq 0.001$; **$p \leq 0.01$; *$p \leq 0.05$. Not significant differences are not shown).

**Figure S2. Spike suppresses IS assembly in CTLs but not in resting $CD8^+$ T cells**. **A**. *Top*, Representative image of a representative $CD8^+$ T cell (day 7) conjugated with SAg-pulsed Raji B cells for 15 min and co-stained with anti-PCNT and anti-GzmB antibodies. The $CD8^+$ T cell was magnified to depict the parameters used for quantification in figures 3, 4, 5, S2 and S3. A line from the center of each lytic granule (marked by GzmB) to the center of the centrosome (marked by PCNT) was drawn to measure the distance (µm) between lytic granules and centrosome. A line from the center of the centrosome (marked by PCNT) of the $CD8^+$ T cell to the center of the contact area with the Raji cell was drawn to measure the distance (µm) between the centrosome and the T cell:APC contact area. *Bottom*, Representative image of a representative $CD8^+$ T cell (day 7) conjugated with SAg-pulsed Raji B cells for 15 min and stained with anti-PTyr mAb. The T cell was magnified to depict the parameters used for quantification in figures 1, 2, 4, 5, S2 and S3. Masks around both the contact area of $CD8^+$ T cell:APC and in the remaining $CD8^+$ T cell area were drawn to measure the mean grey value. The recruitment index (RI) was calculated as specified. **B**. Immunofluorescence analysis of PTyr in CTLs (day 7) pre-treated with either vehicle (PBS) or different concentrations of Spike Wuhan (Spike W), then mixed with Raji cells (APCs) either unpulsed or pulsed with a combination of SEA, SEB and SEE (SAgs), and incubated for 15 min at 37°C. *Left*, Quantification (%) of conjugates harboring PTyr staining at the IS (100 cells/sample, n=3, One-way ANOVA test). *Right*, Relative PTyr fluorescence intensity at the IS (10 cells/sample, n=3, Kruskal-Wallis test). **C-E.** *To*p, Immunofluorescence analysis of PTyr in freshly purified $CD8^+$ T cells (day 0) pre-treated with either vehicle (PBS) or 0.05μg/μl Spike Wuhan (Spike W), then mixed with Raji cells (APCs) either unpulsed or pulsed with a combination of SEA, SEB and SEE (SAgs), and incubated for 15 min at 37°C. The histograms show the quantification (%) of conjugates harboring PTyr (**C**), CD3ζ (**D**) or PCNT (**E**) staining at the IS (≥50 cells/sample, n=3, One-way ANOVA test). *Bottom*, Relative PTyr (**C**) and CD3ζ (**D**) fluorescence intensity at the IS (recruitment index) (10 cells/sample, n=3, Kruskal-Wallis test). Measurement of the distance (μm) of the centrosome (PCNT) from the T cell-APC contact site (**E**) (10 cells/sample, n=3, Kruskal-Wallis test). ****$p \leq 0.0001$; *** $p \leq 0.001$; **$p \leq 0.01$; *$p \leq 0.05$.

**Figure S3. ACE2 suppresses IS assembly in CTLs at an early step. A-C.** *Top*, Quantification (%) of 5-min and 15-min conjugates harboring PTyr (**A**), CD3ζ (**B**) or P-ZAP-70 (**C**) staining at the IS (≥50 cells/sample, n=3, One-way ANOVA test). CTLs (day 7), pre-treated with vehicle (PBS) or 2 μg/ml anti-ACE2 Ab (ACE2), were conjugated with Raji cells (APCs) in the absence or presence of SAgs and either an anti-ACE2 Ab, or angiotensin II (AngII), or the peptide angiotensin 1-7 (Ang 1-7) (≥50 cells/sample, n=3, One-way ANOVA test). *Bottom*, Relative PTyr (**A**), CD3ζ (**B**) or ZAP-70 (**C**) fluorescence intensity at the IS (recruitment index) (10 cells/sample, n=3, Kruskal-Wallis test). **D**. *Left*, Quantification (%) of 5-min and 15-min conjugates harboring PCNT staining at the IS (≥50, n=3, One-way ANOVA test). *Right*, Measurement of the distance (μm) of the centrosome (PCNT) from the T cell-APC contact site (10 cells/sample, n=3, Kruskal-Wallis test). Conjugates were formed as described in panel A. **G**. Quantitative RT-PCR of human MAS1 mRNA in purified $CD8^+$ T cells at days 0, 5 and 7 after stimulation with anti-CD3/CD28 mAb-coated beads in the presence of IL-2 (n=3). ****$p$≤0.0001; *** $p$≤0.001; **$p$≤0.01; *$p$≤0.05. Not significant differences are not shown.

**Videos 1-3**. 3-D reconstruction of a representative CTL-APC conjugate formed with either SAg-unpulsed (video 1) or SAg-pulsed (videos 2 and 3) APCs and co-stained for PTyr (orange) and CD3ζ (green). CTLs used in SAg-specific conjugates were pre-treated with either vehicle (video 2) or Spike-W (video 3).

**Videos 4-6**. 3-D reconstruction of representative CTL-APC conjugate formed with either SAg-unpulsed (video 4) or SAg-pulsed (videos 5 and 6) APCs and co-stained for PNCT (orange) and GzmB (green). CTLs used in SAg-specific conjugates were pre-treated with either vehicle (video 5) or Spike-W (video 6).

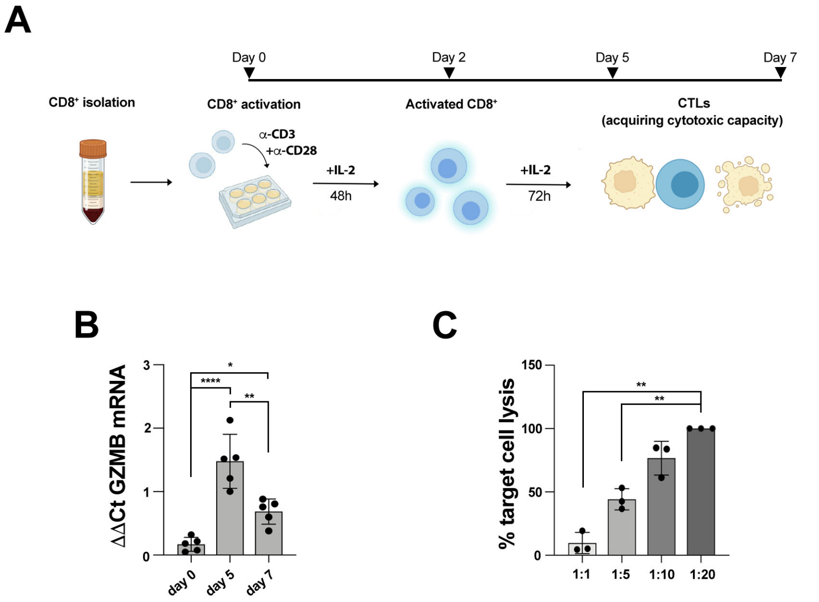
A
Day 0
Day 2
Day 5
Day 7
CD8+ isolation
CD8+ activation
Activated CD8+
CTLs
(acquiring cytotoxic capacity)
α-CD3
+α-CD28
+IL-2
48h
+IL-2
72h
B
ΔΔCt GZMB mRNA
0
1
2
3
*
****
**
day 0
day 5
day 7
C
% target cell lysis
0
50
100
150
**
**
1:1
1:5
1:10
1:20

Figure S1

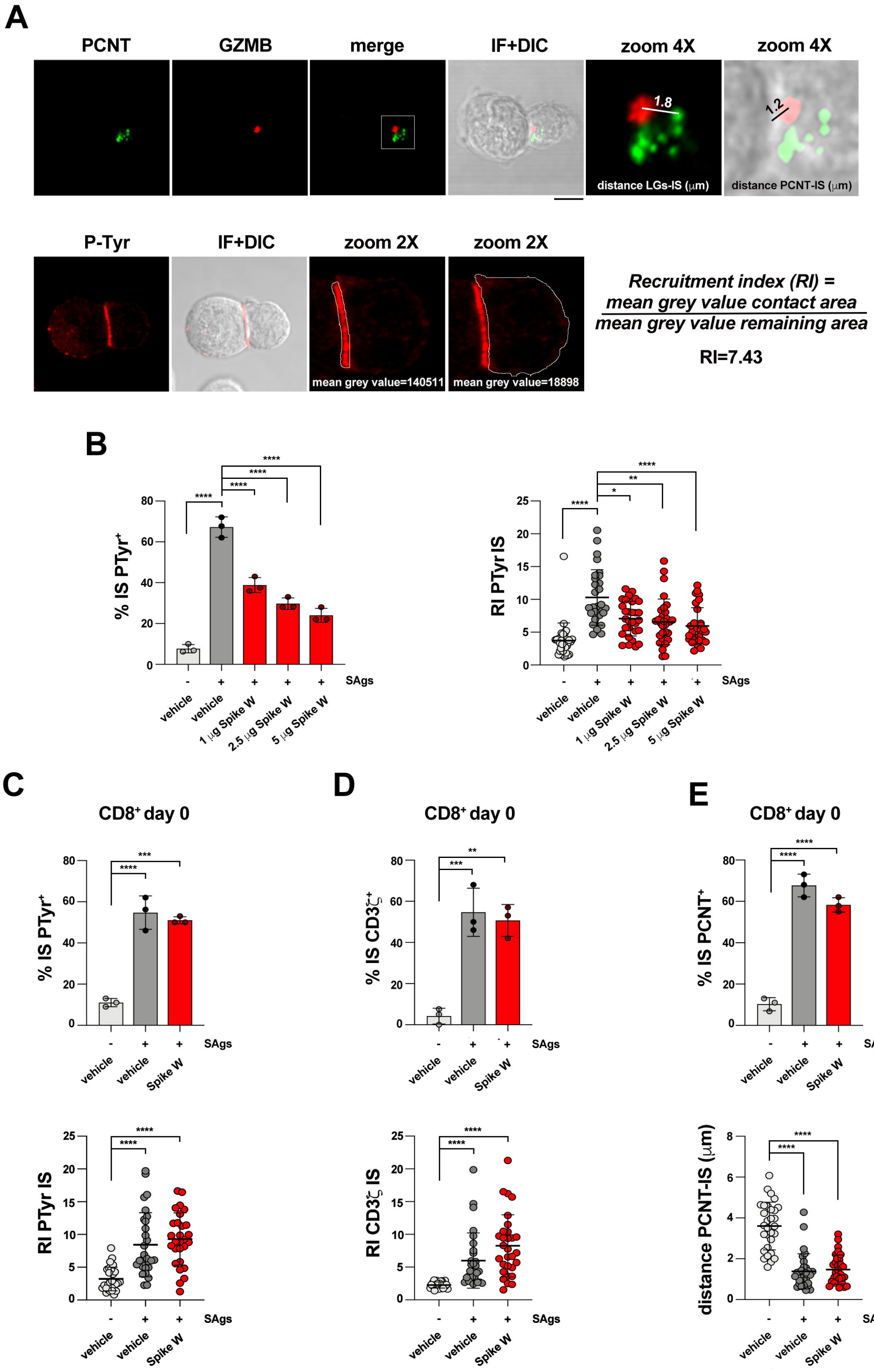

A
PCNT
GZMB
merge
IF+DIC
zoom 4X
zoom 4X
1.8
1.2
distance LGs-IS (μm)
distance PCNT-IS (μm)
P-Tyr
IF+DIC
zoom 2X
zoom 2X
mean grey value=140511
mean grey value=18898
Recruitment index (RI) =
mean grey value contact area
mean grey value remaining area
RI=7.43
B
% IS PTyr+
RI PTyr IS
SAgs
vehicle
vehicle
1 μg Spike W
2.5 μg Spike W
5 μg Spike W
C
CD8+ day 0
% IS PTyr+
RI PTyr IS
D
CD8+ day 0
% IS CD3ζ+
RI CD3ζ IS
E
CD8+ day 0
% IS PCNT+
distance PCNT-IS (μm)
Spike W


Figure S2

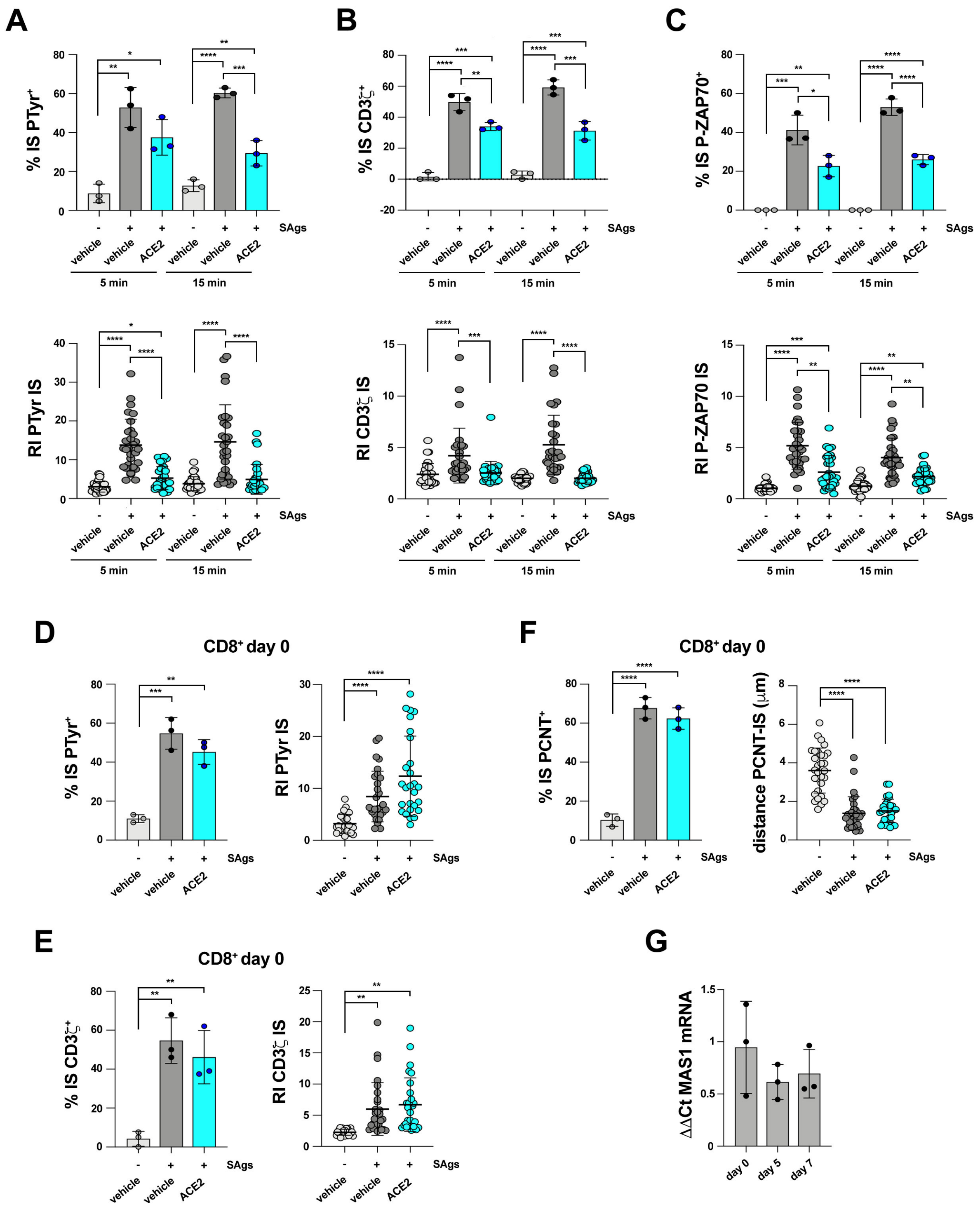
A
% IS PTyr+
RI PTyr IS
B
% IS CD3ζ+
RI CD3ζ IS
C
% IS P-ZAP70+
RI P-ZAP70 IS
SAgs
vehicle
ACE2
5 min
15 min
D
CD8+ day 0
% IS PTyr+
RI PTyr IS
F
CD8+ day 0
% IS PCNT+
distance PCNT-IS (μm)
E
CD8+ day 0
% IS CD3ζ+
RI CD3ζ IS
G
ΔΔCt MAS1 mRNA
day 0
day 5
day 7

**Figure S3**
